# A Comprehensive Enumeration of the Human Proteostasis Network. 2. Components of the Autophagy-Lysosome Pathway

**DOI:** 10.1101/2023.03.22.533675

**Authors:** The Proteostasis Consortium, Overall coordination, Suzanne Elsasser, Lisa P. Elia, Richard I. Morimoto, Evan T. Powers, Harvard Medical School group, Suzanne Elsasser, Daniel Finley, University of California, San Francisco and Gladstone Institutes group I, Beatrice Costa, Maher Budron, Zachary Tokuno, Shijie Wang, Rajshri G. Iyer, Bianca Barth, Eric Mockler, Lisa P. Elia, Steve Finkbeiner, University of California, San Francisco group II, Jason E. Gestwicki, Northwestern University group, Reese A. K. Richardson, Thomas Stoeger, Richard I. Morimoto, The Scripps Research Institute group, Ee Phie Tan, Qiang Xiao, Christian M. Cole, Lynée A. Massey, Dan Garza, Evan T. Powers, Jeffery W. Kelly, Stanford University group, T. Kelly Rainbolt, Ching-Chieh Chou, Vincent B. Masto, Judith Frydman, New York University group, Ralph A. Nixon

## Abstract

The condition of having a healthy, functional proteome is known as protein homeostasis, or proteostasis. Establishing and maintaining proteostasis is the province of the proteostasis network, approximately 2,700 components that regulate protein synthesis, folding, localization, and degradation. The proteostasis network is a fundamental entity in biology that is essential for cellular health and has direct relevance to many diseases of protein conformation. However, it is not well defined or annotated, which hinders its functional characterization in health and disease. In this series of manuscripts, we aim to operationally define the human proteostasis network by providing a comprehensive, annotated list of its components. We provided in a previous manuscript a list of chaperones and folding enzymes as well as the components that make up the machineries for protein synthesis, protein trafficking into and out of organelles, and organelle-specific degradation pathways. Here, we provide a curated list of 838 unique high-confidence components of the autophagy-lysosome pathway, one of the two major protein degradation systems in human cells.

## Introduction

Proteostasis is the maintenance of the proteome in a healthy, functional state.^1^ Proteostasis is enabled by the proteostasis network, a collection of cellular components responsible for managing the synthesis, folding, trafficking, and degradation of proteins.^1–5^ Although the term “proteostasis network” was introduced over a decade ago,^1^ the network has remained poorly defined as there is no comprehensive accounting of its components. This problem is especially acute considering the important role of proteostasis in aging-related neurodegenerative diseases.^6–11^ To address this problem, we have been creating a detailed enumeration of the proteostasis network. The first installment of this list was reported recently and included the machinery of protein synthesis, chaperones, folding enzymes, the systems for trafficking proteins into and out of organelles, and organelle-specific degradation pathways.^12^ Here we report the next installment, a comprehensive list of the components of the autophagy-lysosome pathway (ALP). The ALP is one of two major systems for the degradation of proteins (the other being the ubiquitin-proteasome system, or UPS, which is the subject of an upcoming manuscript). In the ALP, the substrates to be degraded are surrounded by a double-membrane vesicle called the autophagosome, which is transported to and fuses with the lysosome for degradation.^13–19^ Within the proteostasis network, the ALP is of particular interest because mutations in many of its components have been found to increase the risk of diseases of failed proteostasis, like Alzheimer’s disease, frontotemporal dementia, Parkinson’s disease, and amyotrophic lateral sclerosis, among others.^11,20^

## Results

### Taxonomic system for annotating proteostasis network components, and criteria for inclusion

We previously introduced a taxonomic annotation system for the proteostasis network.^12^ This system was constructed to convey at a glance a component’s role in proteostasis. It consists of five levels: Branch, Class, Group, Type, and Subtype. The Branch annotation is the broadest and refers to a component’s localization or membership in an overarching pathway. There are eight Branches of the proteostasis network (PN; Figure 1), six of which we described previously (cytonuclear proteostasis, ER proteostasis, mitochondrial proteostasis, nuclear proteostasis, cytosolic translation, and proteostasis network regulation).^12^ The ALP is the seventh Branch of the proteostasis network, and the UPS is the eighth. The Class, Group, Type and Subtype annotations give increasingly specific descriptions of a component’s role in proteostasis. We endeavored to use as few descriptors as possible for each component. Thus, many components lack Type and Subtype annotations. Also, some components have multiple roles in proteostasis and therefore have multiple sets of annotations.

**Figure 1.**
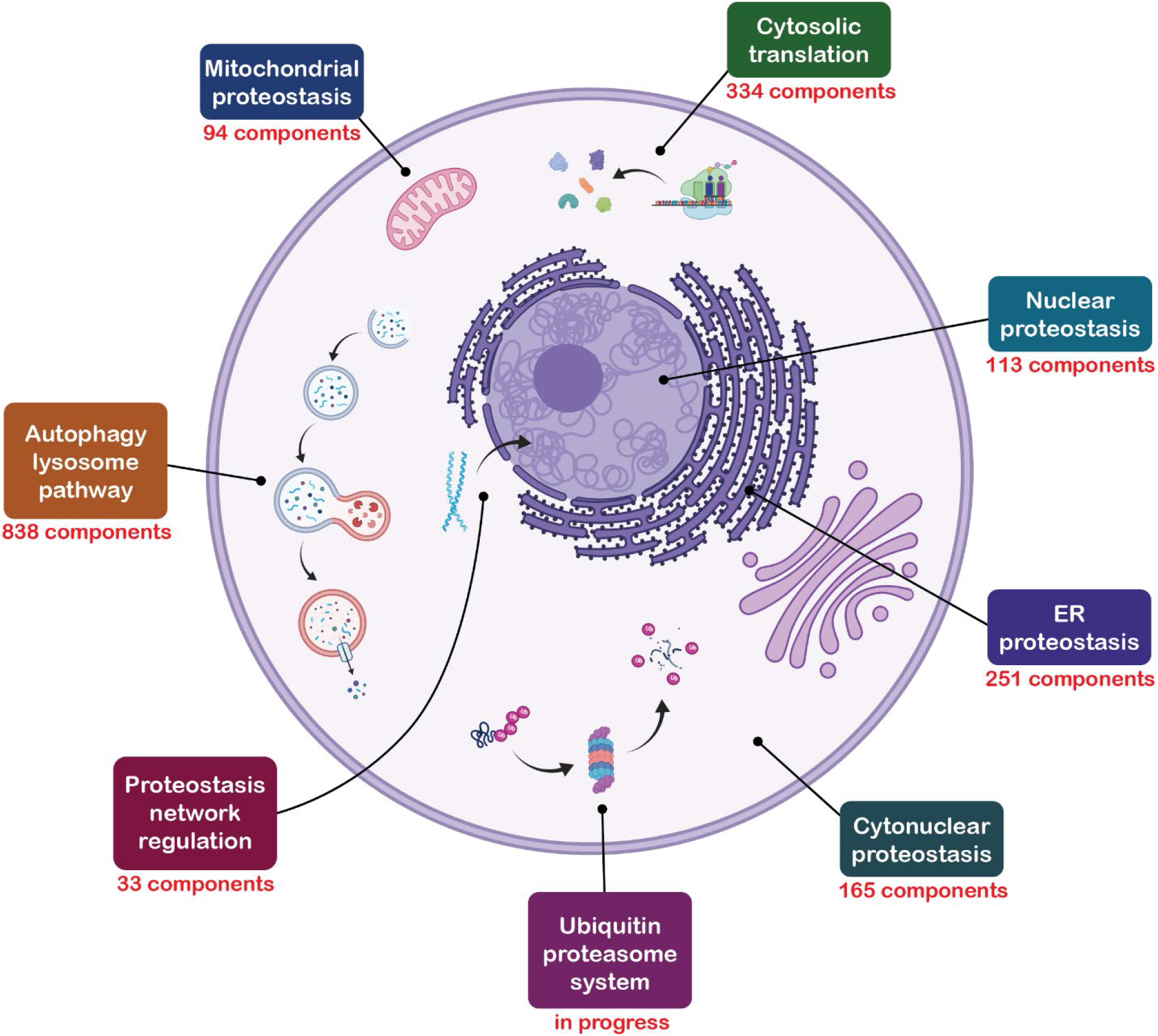
The proteostasis network. The branches of the proteostasis network are called out in boxes. The number of unique components is shown for the ALP and the six branches described in our previous manuscript, from which this figure is adapted.^12^ Parts of this figure were created with BioRender.com
.

We previously described two criteria that we employ to decide whether components should be included in the proteostasis network.^12^ The first is the “entity-based” criterion, in which entities (proteins, protein complexes, non-coding RNA, etc.) are included because there is experimental evidence for their having a role in proteostasis. The second is the “domain-based” criterion, in which components are included because they have at least one structural domain that is characteristic of protein families known to be involved in proteostasis. We primarily used the entity-based criterion applied to existing lists of autophagy components (for example, autophagy-related Gene Ontology annotations,^21,22^ the KEGG pathway “Autophagy - Animal”,^23^ and previous compilations of ALP components,^24–26^ especially the recent list by Bordi and co-workers^27^) to populate our initial list of ALP components. This initial list was then expanded through a comprehensive literature search as described in the Methods section. The domain-based criterion was also used in a few cases. For example, MAP1LC3B2 was included because it is a homolog of the yeast protein ATG8.^28,29^ ATG8 is central to the yeast ALP^30,31^ and six of its seven human homologs have also been shown to function in the ALP, but to our knowledge MAP1LC3B2 has not.^28^

While the scope of the PN naturally encompasses the fate of proteins, our enumeration extends slightly beyond the bounds of the protein world. For example, our description of selective autophagy includes all forms of it, including those by which biomolecules other than proteins are degraded, such as lipophagy.^32^ We also included all the catabolic enzymes of the lysosome in the ALP, not just those that catalyze the cleavage of peptide bonds, as well as those components involved in maintaining the acidic lysosomal environment. However, we did not include components that are generally involved in lysosomal or endosomal trafficking. We also did not include components associated only with non-canonical autophagy or with secretory autophagy.^33–35^ These pathways may be included in later versions of the ALP when their significance and mechanisms are better understood.

### Application of the taxonomic scheme to the ALP

Our taxonomy of the ALP is underpinned by the temporal progression of autophagy, much of which was first elucidated by Ohsumi and co-workers.^36^ Autophagy is governed by signaling pathways that regulate flux through the ALP. These inputs—including, for example, nutrient, hormone, energy, and stress signals—are mostly communicated through mTORC1 (mammalian target of rapamycin complex 1) and AMPK (AMP-activated protein kinase), which are regulators of numerous key pathways in addition to autophagy, like cell metabolism and energy homeostasis.^37–42^ These kinases modulate autophagy through modification of the ULK complex (unc-51 like autophagy activating kinase), with mTORC1 and AMPK exerting inhibitory and activating effects, respectively, through phosphorylation at specific sites (for example, Ser757 of ULK1 by mTORC1 and Ser317/Ser777 of ULK1 by AMPK).^38,41^ When the ULK complex is activated it is recruited to the site where the autophagophore (as the nascent autophagosome is known) formation is initiated; sometimes, this is at a structure in the ER membrane known as the omegasome.^43^ There, the ULK complex activates the PI3KC3 complex 1 (class III phosphatidylinositol 3-kinase)^44^ which phosphorylates phosphatidylinositol (PI) at the 3 position to yield phosphatidylinositol 3-phosphate (PI(3)P).^14,45–47^ PI(3)P recruits the ATG2-WIPI complex (ATG = autophagy related; WIPI = WD repeat domain, phosphoinositide interacting), which rapidly transports lipids to the growing autophagophore.^14,48,49^ The ATG2-WIPI complex also recruits the ATG5-ATG12-ATG16 complex,^50^ which catalyzes the conjugation of phosphatidylethanolamine to human ATG8 orthologs like the MAP1LC3 or GABARAP proteins on the autophagophore membrane (MAP1LC3 = microtubule associated protein light chain 3; GABARAP = GABA type A receptor-associated protein).^51–53^ These lipidated proteins are critical components of the autophagosome,^28,54–57^ especially for substrate recognition.^58–61^

The autophagophore grows until it surrounds its substrate, whether that is a portion of cytosol, as in bulk macroautophagy, or other substrates such as protein aggregates or entire organelles, as in selective autophagy. The substrates of selective autophagy are bound by receptors that recognize substrates by various mechanisms (often involving ubiquitin^59^), and recognize autophagosomes via their displayed ATG8 homologs.^58,60,61^ The autophagophore is sealed with the assistance of ESCRT complexes (endosomal complexes required for transport), yielding an autophagosome.^14,62^ Autophagosomes are transported to through the cell^19,63^ to fuse with lysosomes, creating autolysosomes,^15^ in which the cargo from the autophagosome is degraded by lysosomal enzymes.^64^ Lysosomes are then regenerated from the autolysosomes by the process of autophagic lysosome reformation.^65^

The Classes in the ALP Branch of the proteostasis network largely reflect the stages described above in the autophagy process and are illustrated in the schematic overview of the ALP in Figure 2. These Classes are “pre-initiation autophagy signaling”; “autophagophore initiation and elongation”; “autophagy substrate selection”; “autophagosome closure, maturation, and lysosome fusion”; “lysosomal catabolism”; and “autophagic lysosome reformation”. The remaining classes are “autophagy gene expression”, which comprises the transcription factors and transcriptional and translational regulators that control the expression of autophagy genes; “chaperone directed autophagy”, which includes components involved in chaperone-mediated autophagy^66^ and chaperone assisted selective autophagy^67^; and “specific function in autophagy unknown”, a small Class that contains components that clearly affect autophagy but through mechanisms that are not yet known. The Groups, Types, and Subtypes further specify a component’s function within the processes denoted by their Class.

**Figure 2.**
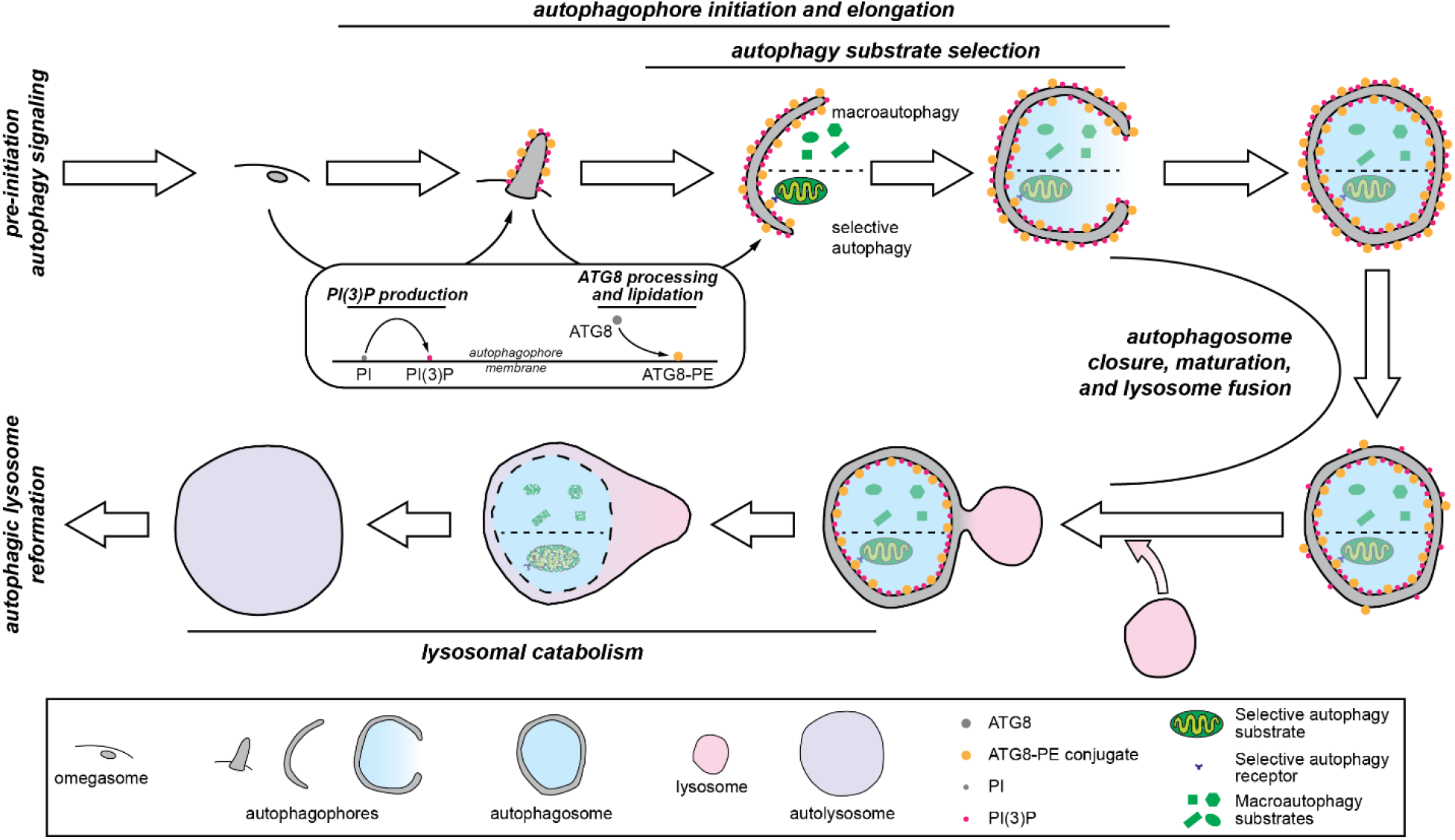
A schematic overview of the ALP. The temporal stages that correspond to Classes in the ALP are highlighted. **Pre-initiation autophagy signaling**: Flux through the ALP is regulated by signaling pathways that are upstream of autophagy initiation. **Autophagophore initiation and elongation**: Once autophagy is initiated, the autophagophore double membrane begins to grow. Initiation sometimes occurs at structures in the ER membrane known as omegasomes, but there are likely many mechanisms by which autophagophores can be nucleated. The growing autophagophore is decorated with PI(3)P (phosphatidylinositol 3-phosphate) and ATG8-PE (ATG8-phosphatidylethanolamine conjugate). **Autophagy substrate selection**: As the autophagophore grows, it encloses whatever components of the cytosol are in its vicinity (in macroautophagy) or components that are recruited via interactions between ATG8 and autophagy receptors (in selective autophagy). **Autophagosome closure, maturation, and lysosome fusion**: When it is large enough, the autophagophore membrane closes to form the autophagosome. As the autophagosome is transported in the cell, its membrane composition is modified in the process of autophagosome maturation, which readies it for fusion with the lysosome to form the autolysosome. **Lysosomal catabolism**: After lysosomal fusion, the autophagosome’s interior membrane and the cargo it contains are digested by lysosomal enzymes. **Autophagic lysosome reformation:** After lysosomal catabolism is complete, the components of the autolysosome are recycled to form new lysosomes.

The application of our taxonomic scheme to an ALP component is illustrated in Figure 3 for ATG4A, a cysteine protease that trims the C-terminal Arg from ATG8 homologs in preparation for their conjugation to phosphatidylethanolamine.^68,69^ The following annotations were assigned to ATG4A: Branch = “autophagy-lysosome pathway”; Class = “autophagophore initiation and elongation”; Group = “ATG8 homolog processing, direct”; Type = “preparation of ATG8 homologs for lipidation”; Subtype = “peptidase that removes C-terminal Arg from ATG8”. It is important to note that these annotations are not unique, as they are shared with ATG4A’s paralogs in the human genome: ATG4B, ATG4C, and ATG4D.^69^ Also, like many other ALP components, ATG4A has two distinct sets of annotations because in addition to its role in preparing ATG8 homologs for lipidation, it is also responsible for delipidating ATG8 homologs and removing them from the autophagosome membrane as the autophagosome matures.^68,70^ The annotation for this role of ATG4A in the ALP is: Branch = “autophagy-lysosome pathway”; Class = “autophagosome closure, maturation, and lysosome fusion”; Group = “regulation of autophagosome membrane composition”; Type = “ATG8 homolog de-lipidation”; Subtype = “releases ATG8 homologs from maturing autophagosome”.

**Figure 3.**
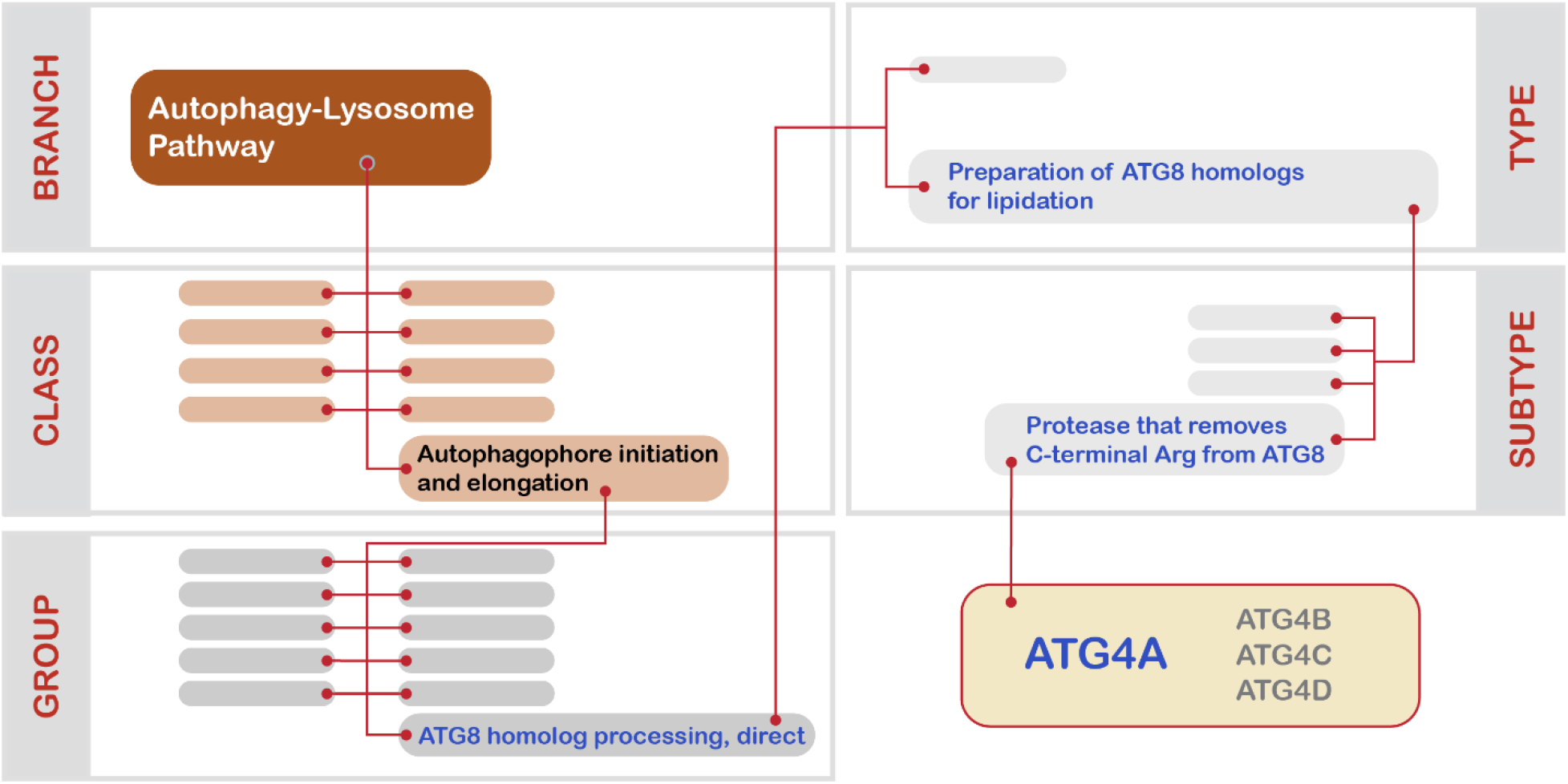
Illustration of the five-level taxonomic scheme applied to proteostasis network components herein using the cysteine protease ATG4A. Blank boxes correspond to categories within each taxonomic level to which ATG4A does not belong. ATG4A has two annotations (see text), but here we illustrate only ATG4A’s annotation in the “autophagophore initiation and elongation” Class.

### The autophagy-lysosome pathway consists of at least 838 unique protein-coding genes

The list of components of the ALP is presented in Supplemental Table 1 (see the Methods section for details on how the list was constructed). There are 985 entries in our list of ALP components, representing 838 unique components. Of these, 714 have a single entry, 108 have two, 11 have three, 4 have four, and 1 (SQSTM1, or sequestosome 1) has six. All the entries have been assigned to one of the nine Classes of the ALP Branch, and all but three have one of the 76 Group annotations. The three exceptions are those components in the “specific function in autophagy unknown” Class. Most of the entries have one of the 138 Type annotations (899 out of 985), but fewer than half of the entries have one of the 78 Subtype annotations (365 out of 985).

To illustrate how the finer-grained annotations were assigned in the ALP, we show how the components of the Class “autophagosome closure, maturation, and lysosome fusion” are distributed within this Class’s various Groups, Types, and Subtypes in Table 1.

**Table 1.**
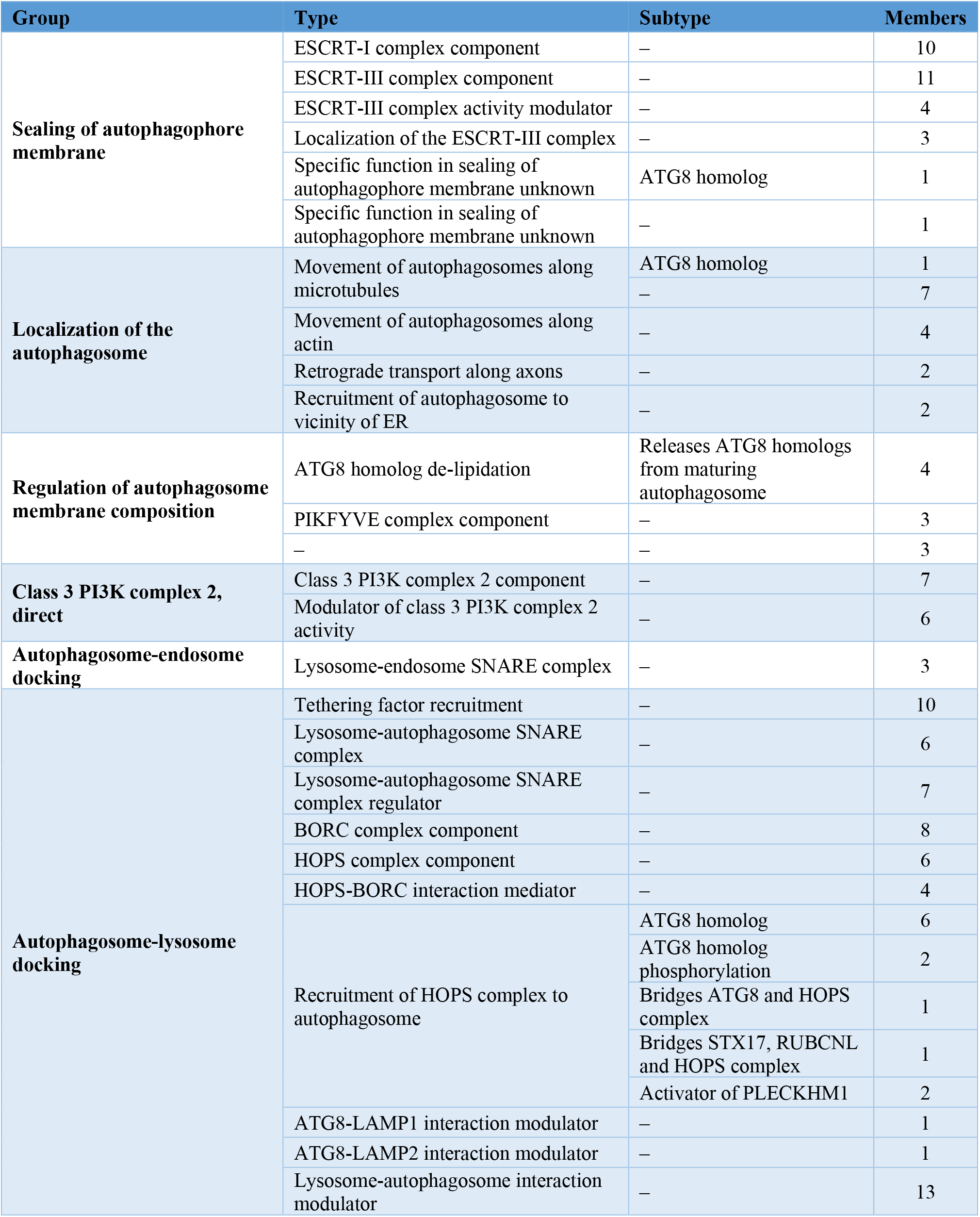
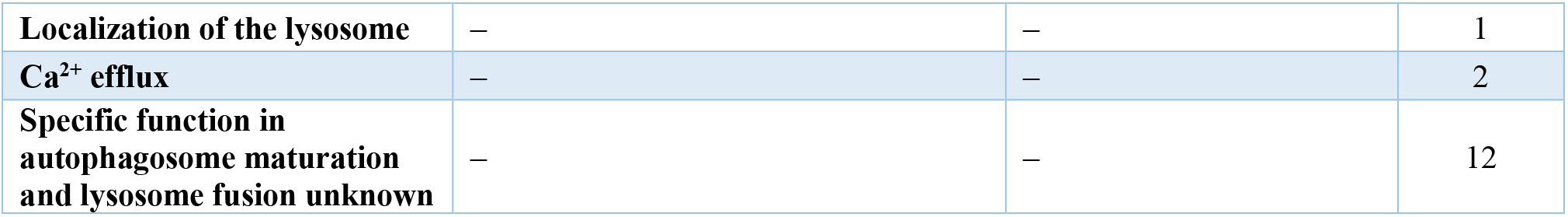
Groups, Types and Subtypes within the “autophagosome closure, maturation, and lysosome fusion” Class.

## Discussion

The ALP is much larger than any of the proteostasis network branches described in our preceding manuscript, consisting of over 800 components. The size of the ALP can be attributed both to the complexity of the physical processes involved—membrane nucleation and growth, substrate selection, vesicle trafficking and fusion—and to the extensive regulation of these processes. This regulation happens at all points in the ALP, but mTORC1,^37,40^ which regulates autophagy initiation, and PI3KC3 complex 1,^71,72^ which regulates the PI(3)P content of the autophagophore, are especially dense nodes for regulatory activity. This level of evolutionary investment into the control systems of the ALP shows how critical it is to tune the level of catabolism to the state of the cell. The significance of the ALP to cellular health is also manifested in the outsized representation of ALP mutations in many aging-related neurodegenerative diseases, as noted above.^11,20^ We expect that the list of ALP components described herein will enable a deeper understanding of the ways that failures of the ALP contribute to the pathogenesis of such diseases by enabling those studying them to see the full array of connections between, say, the results obtained from patient-derived multiomic datasets and the ALP. In this way, new targets could be discovered for diseases with known links to the ALP. Moreover, our comprehensive accounting of ALP components could reveal new roles for the ALP in diseases that are not currently associated with it, enabling the exploration of new therapeutic strategies.

Finally, we note that we view the list of annotations presented herein as a first version. We will regularly update our list of ALP components as new information becomes available. Suggestions for components that should be included or removed, or annotations that should be added, deleted, or changed can be sent to.

## Methods

Our “entity-based” and “domain-based” criteria for inclusion of components in the proteostasis network were described in the previous manuscript in this series.^12^ To generate the initial lists of ALP components we relied on the existing autophagy-related Gene Ontology annotations^21,22^ the KEGG pathway “Autophagy - Animal”,^23^ and previous compilations of ALP components^24–27^ supplemented with recent reviews of either autophagy in general or specific aspects of autophagy.^11,13–20,28,37,58–61,63–66,73–76^ The initial list consisted of ~730 components. We then explored the literature for genes with associations with autophagy that were either newly discovered or not recognized in the sources referenced above. We used the Gene database from NCBI^77^ to find papers related to each gene in the genome that had been published as of June 2021. The titles, abstracts, and MeSH terms for these papers were downloaded and scanned for the word fragment “-autopha-” to identify papers that could conceivably report a role for the gene of interest in autophagy. We selected ~420 genes for further evaluation that had four or more related papers in which the fragment “-autopha-” occurred and were not already in our preliminary list. The relevant abstracts (and the manuscripts themselves, as necessary) for each of these genes were read by members of the Consortium to determine if they should be considered as candidates for inclusion in the ALP. Candidate genes were proposed to the Consortium’s PN annotation subgroup and inclusion and exclusion decisions were made collectively by the subgroup. Approximately 110 components were added to the preliminary list through this process. Each ALP component entry in Supplemental Table 1 has a note justifying its inclusion in the ALP and explaining the annotation, as well as links to supporting literature, in the last few columns of the table.

## Acknowledgements

We gratefully acknowledge funding from the National Institutes of Health (National Institute on Aging P01AG054407 to R. I. M., D. F., S. F., J. E. G., E. T. P., J. W. K., and J. F.; P01AG017617 and R01AG062376 to R. A. N.; K99AG068544 to T. S.; and K99AG080109 to E. P. T.)

